# The plant-specific SCL30a SR protein regulates ABA-dependent seed traits and salt stress tolerance during germination

**DOI:** 10.1101/2021.10.13.464208

**Authors:** Tom Laloum, Sofia D. Carvalho, Guiomar Martín, Dale N. Richardson, Tiago M. D. Cruz, Raquel F. Carvalho, Kevin L. Stecca, Anthony J. Kinney, Mathias Zeidler, Inês C. R. Barbosa, Paula Duque

**Affiliations:** Instituto Gulbenkian de Ciência, Rua da Quinta Grande, 6, 2780-156 Oeiras, Portugal; Crop Genetics Research and Development, DuPont Experimental Station, Wilmington, Delaware 19880-0353, USA; Institute of Plant Physiology, Justus-Liebig-University Gießen, Senckenbergstraße 3, D-35390 Gießen, Germany

## Abstract

SR (serine/arginine-rich) proteins are conserved RNA-binding proteins best known as key regulators of splicing, which have also been implicated in other steps of gene expression. Despite mounting evidence for their role in plant development and stress responses, the molecular pathways underlying SR protein regulation of these processes remain elusive. Here we show that the plant-specific SCL30a SR protein negatively regulates abscisic acid (ABA) signaling to control important seed traits and salt stress responses during germination in Arabidopsis. The *SCL30a* gene is upregulated during seed imbibition and germination, and its loss of function results in smaller seeds displaying enhanced dormancy and elevated expression of ABA-responsive genes as well as of genes repressed during the germination process. Moreover, the knockout mutant is hypersensitive to ABA and high salinity, while transgenic plants overexpressing *SCL30a* exhibit reduced ABA sensitivity and enhanced tolerance to salt stress during seed germination. An ABA biosynthesis inhibitor rescues the mutant’s enhanced sensitivity to stress, and epistatic analyses confirm that this hypersensitivity requires a functional ABA pathway. Finally, seed ABA levels are unchanged by altered *SCL30a* expression, indicating that the SR protein positively regulates stress tolerance during seed germination by reducing sensitivity to the phytohormone. Our results reveal a new key player in ABA-mediated control of early development and stress response, and underscore the role of plant SR proteins as important regulators of the ABA signaling pathway.

**Author Summary:** Seed germination is a critical step in plant development determining the transition to aerial growth and exposure to a more challenging environment. As such, seeds have evolved mechanisms that prevent germination under adverse conditions, thereby increasing the chances of plant survival. As a general regulator of plant development and a key mediator of stress responses, the hormone abscisic acid (ABA) promotes a prolonged non-germinating state called dormancy, influences seed size and represses germination under environmental stress. Here, we show that an RNA-binding protein, SCL30a, controls seed size, dormancy, germination and tolerance to high salinity in the model plant *Arabidopsis thaliana*. Loss of *SCL30a* gene function results in smaller and more dormant seeds with reduced ability to germinate in a high-salt environment; by contrast, *SCL30a* overexpression produces larger seeds that germinate faster under salt stress. Using a large-scale gene expression analysis, we identify the ABA hormonal pathway as a putative target of SCL30a. We then use genetic and pharmacological tools to unequivocally demonstrate that the uncovered biological functions of SCL30a are achieved through modulation of the ABA pathway. Our study reveals a novel regulator of key seed traits and has biotechnological implications for crop improvement under adverse environments.

## Introduction

Seed germination begins with rehydration (imbibition) and expansion of the embryo by cell elongation, which leads to rupture of the weakened seed coat and emergence of the radicle [1]. During water uptake, a prolonged non-germinating state termed seed dormancy must be relieved before protrusion of the radicle [2]. The completion of seed germination marks a key developmental milestone in the life cycle of higher plants, being essential for the establishment of a viable plant. The germination process is highly regulated by both endogenous and environmental signals that determine the dormancy status of the seed and its aptitude to germinate [3].

The plant hormone abscisic acid (ABA) promotes seed maturation and dormancy while inhibiting seed germination, thus acting as a key regulator of this critical developmental step [4,5]. In fact, mutations that affect components of ABA biosynthesis (e.g., *aba2*) or signaling (e.g., *snrk2.2/3/6*) exhibit reduced seed dormancy and precocious germination [6,7]. ABA has also more recently been implicated in the control the seed’s final size, with ABA production and signaling modulating the expression of *SHB1*, a main regulator of endosperm cellularization during seed development [8].

Apart from regulating key developmental processes such as seed germination, ABA is a known major mediator of osmotic stress responses, also in seeds where it acts as an integrator of different environmental signals to repress germination under unfavorable conditions [5]. While numerous studies have deciphered the genetic components and transcriptional mechanisms underlying seed germination and osmotic stress responses, the involvement of posttranscriptional gene regulation, namely of alternative splicing, is beginning to unfold [9–11].

RNA splicing, which excises introns from the precursor mRNA (pre-mRNA) and joins the flanking exonic sequences to generate mature transcripts, is an essential step in eukaryotic gene expression. This process involves the recognition of intronic sequences called splice sites by the spliceosome, a large molecular complex consisting of five small nuclear ribonucleoproteins (snRNPs) and numerous spliceosome-associated proteins that assemble at introns in a precise order [12,13]. The differential recognition of splice sites results in alternative splicing, which allows a single gene to express multiple mRNA variants and hence greatly contributes to transcriptome diversification.

SR (serine/arginine-rich) proteins are multi-domain, non-snRNP spliceosomal factors that regulate pre-mRNA splicing. These RNA-binding proteins use one or two of their N-terminal RNA recognition motifs (RRMs) to bind to specific cis-acting elements in pre-mRNAs and enhance or repress splicing [14]. SR proteins recruit core spliceosomal factors to pre-mRNAs through their C-terminal arginine/serine (RS) domain, which acts a protein-protein interaction module [15]. The RS domain is also subjected to numerous reversible phosphorylation events that control SR protein activity and subcellular localization [16,17].

Apart from pre-mRNA splicing, non-canonical functions for SR proteins in pre-and post-splicing activities have been emerging, highlighting their multifaceted roles as important coordinators of nuclear and cytoplasmic gene expression machineries [18,19]. In one example, the mammalian SR protein SRSF2 was shown to mediate the activation of the paused Pol II by releasing the positive transcription elongation factor b (p-TEFb) from inhibitory 7SK ribonucleoprotein complexes, thus promoting transcriptional elongation [20]. Furthermore, changes in SRSF2 levels have been shown to affect the accumulation of Pol II at gene loci [21]. More generally, SR proteins influence gene transcription by directly or indirectly interacting with the C-terminal domain of RNA Pol II during their assembly as RNA processing factors [18]. Animal SR proteins have also been shown to influence mRNA export, translation and decay by interacting with major components of the molecular complexes regulating these processes [19].

Functional analyses of individual SR and SR-like proteins in plants have identified specific roles for these proteins in stress and ABA responses. The Arabidopsis RS40 and RS41 were found to interact in nuclear speckles with HOS5, a KH-domain RNA-binding protein, and FRY2/CPL1, a major player in the co-transcriptional processing of nascent transcripts, with knockout mutants of these two SR proteins displaying hypersensitivity to ABA during seed germination as well as to the inhibitory effect of salt on root elongation [22]. RSZ22 is a putative dephosphorylation target of the Clade A protein phosphatase 2C HAI1, a major component of ABA and osmotic stress signaling in Arabidopsis [23]. The SR-like SR45a was recently shown to inhibit salt stress tolerance in Arabidopsis by interacting with the RNA cap-binding protein CBP20 and regulating alternative splicing of transcripts involved in the response to high salinity [24]. In addition to salt stress responses [25], the other Arabidopsis SR-like protein, SR45, regulates sugar responses by repressing both ABA signaling and glucose-induced accumulation of the hormone [26,27], with SR45-bound transcripts being markedly enriched in ABA signaling functions [28]. In support of a conserved role for these proteins in splicing regulation, SR45 and SR45a interact with the spliceosomal components U1-70K and U2AF35b [24,29] involved in the recognition of 5’ and 3’ splice sites, respectively.

The Arabidopsis genome encodes 18 SR proteins, 10 of which are orthologs of the human ASF/SF2, 9G8 or SC35, while members of the RS, RS2Z and SCL subfamilies are plant-specific [30]. SCL30a belongs to the latter subfamily, whose four members are similar to SC35 but display a distinctive short N-terminal charged extension rich in arginines, serines, glycines and tyrosines [30]. SCL30a interacts with the RS2Z33 SR protein [31] and acts redundantly with its paralog SCL33 to control alternative splicing of a specific intron in the SCL33 pre-mRNA [32]. A more recent study described pleiotropic developmental phenotypes for a quintuple mutant of the four SCL and the SC35 genes, including serrated leaves, late flowering, shorter roots and abnormal silique phyllotaxy, while the corresponding single mutants did not show obvious phenotypic alterations [33]. Furthermore, all four SCL members (SCL28, SCL30, SCL30a and SCL33) and SC35 localize in nuclear speckles and interact with major components of the early spliceosome machinery U170K and U2AF65a [33], corroborating their function as splicing regulators. Interestingly, these five Arabidopsis SR proteins were also recently reported to interact with the NRPB4 subunit of RNA Pol II, indicating a potential role in the regulation of transcription [33].

Here, we characterized the plant-specific *SCL30a* gene in Arabidopsis and found that the encoded protein regulates seed size, dormancy and germination. Loss of *SCL30a* function affects alternative splicing of a limited number of genes, but upregulates expression of many osmotic stress and ABA-responsive genes. In agreement with this, the *scl30a-1* loss-of-function mutant displays strong hypersensitivity to ABA and salt stress during seed germination. Conversely, overexpression of *SCL30a* reduces ABA sensitivity and confers seed tolerance to salt stress during germination. Epistatic and pharmacological analyses demonstrate that SCL30a’s function in seeds and salt stress tolerance depends on ABA synthesis and signaling, demonstrating a key role for this RNA-binding protein in ABA-mediated responses.

## Results

### *SCL30a* expression is markedly induced during seed germination

To initiate the characterization of the Arabidopsis *SCL30a* gene and investigate its expression pattern, we generated transgenic plants expressing the β-glucuronidase (*GUS*) reporter gene under the control of the *SCL30a* endogenous promoter. The *SCL30a* promoter was active throughout plant development (Fig 1A). We observed GUS staining in vascular tissues and actively dividing cells, such as in the shoot meristem and young leaves (Fig 1A (a)), the primary root tip (Fig 1A (b)) and lateral root primordia (Fig 1A (c)). At the reproductive phase, *SCL30a* appeared to be particularly expressed in the pistil tip, the vasculature tissue of sepals, the stamen filaments and pollen grains (Fig 1A (d)) of developing flowers. In embryonic tissues, the *SCL30a* promoter was active from the early — globular and heart (Fig 1A (e-g)) — to the late — torpedo and mature embryo (Figure 1A (h-j)) — stages of embryo development. Finally, in imbibed mature seeds, GUS staining was detected in the whole embryo (Fig 1A (k)) as well as strongly in the seed coat (Fig 1A (l)), but is mainly expressed at the radicle tip during germination (Fig 1A (m)).

**Fig 1.**
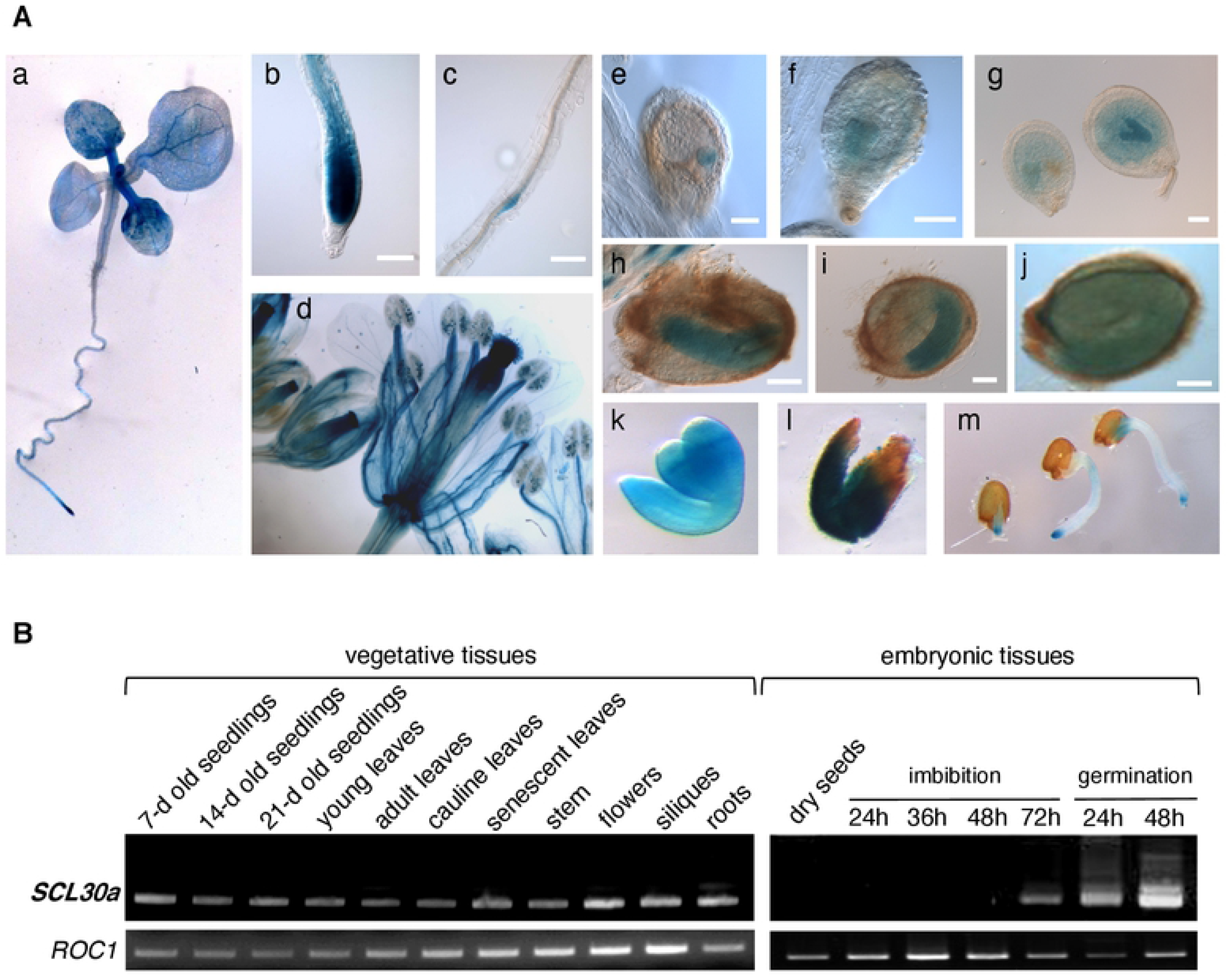
*SCL30a* promoter activity and expression pattern in Arabidopsis. **(A)** Differential interference contrast microscopy images of GUS-stained transgenic plants carrying the *promSCL30a:GUS* reporter construct. *SCL30a* promoter activity in 2-week-old seedlings (a), the primary root tip (b), a lateral root primordium (c), mature and immature flowers (d), developing embryos (e-j), the embryo (k) and testa (l) from imbibed mature seeds, and seeds germinated for 1-2 days (m). Scale bars, 100 μm. **(B)** RT-PCR analysis of *SCL30a* transcript levels in vegetative and embryonic tissues of wild-type (Col-0) plants. The location of the F1 and R1 primers is shown in S1A Fig. Expression of the *cyclophilin* (*ROC1*) gene was used as a loading control.

In parallel, we used RT-PCR to study the development- and tissue-specific expression pattern of *SCL30a*. Consistent with the established *promoter:GUS* expression profile, *SCL30a* was expressed both in young seedlings and at later developmental stages, with its mRNA being detected in different aerial tissues, such as leaves, stem, flowers and siliques, but also in roots (Fig 1B). In embryonic tissues, although *SCL30a* transcripts were undetectable in dry seeds, gene expression was clearly observed at 3 days of seed imbibition at 4 °C and increased sharply during the first hours of germination upon transfer to 22 °C and light (Fig 1B).

Both animal and plant pre-mRNAs encoding splicing components appear to be particularly prone to alternative splicing themselves. This has been shown to lead to a dramatic increase of the transcriptome complexity of the Arabidopsis SR protein family [34,35], prompting us to examine the splicing pattern of the *SCL30a* gene. Although only one transcript has been annotated (https://www.arabidopsis.org), cloning and sequencing of the PCR products amplified from the *SCL30a* cDNA identified three alternative mRNAs (S1 Fig), consistent with the information available in PASTDB (http://pastdb.crg.eu), a recently developed transcriptome-wide resource of alternative splicing profiles in Arabidopsis [36]. The shortest and by far most expressed *SCL30a.1* transcript (Fig 1B and S1B Fig) is predicted to encode the full-length protein, while the other two splice variants encode putative severely truncated proteins (S1A Fig).

Thus, the Arabidopsis *SCL30a* gene, which produces at least three alternative transcripts, displays ubiquitous expression in vegetative tissues and is induced during seed germination.

### Loss of *SCL30a* function reduces seed size, enhances seed dormancy and delays germination

To investigate the biological roles of *SCL30a*, we isolated a homozygous T-DNA mutant line, SALK_041849, carrying the insertion in the gene’s third exon (S1A Fig). RT-PCR analysis of *SCL30a* expression in this *scl30a-1* mutant using primers annealing upstream of the insertion site revealed transcript levels comparable to the Col-0 wild type, but no expression was detected when primers flanking or annealing downstream of the T-DNA were used (S1B Fig). Consistent with the location of the insertion, no splice variants were detected in the mutant, which only expresses a truncated *SCL30a* transcript lacking the sequence corresponding to the entire RS domain as well as most of the RRM (S1 Fig). These results indicate that the *scl30a-1* allele is a true loss-of-function mutant.

Given that we observed no notable defects in adult plants and the marked induction of the *SCL30a* gene during seed imbibition and germination (see Fig 1), we focused our phenotypical analysis of the *scl30a-1* mutant on embryonic tissues. Notably, mature *scl30a-1* seeds displayed a significant reduction in size, with dry and imbibed mutant seeds being 12 % and 14 % smaller, respectively, than seeds of wild-type plants (Fig 2A). Correlating with their smaller size, dry mature *scl30a-1* seeds showed reduced weight when compared to the wild type, but no significant changes in their relative moisture, protein or oil content (S1 Table). A more detailed compositional analysis revealed only minor changes in the relative levels of a few unsaturated fatty acids and the trisaccharide raffinose (S1 Table), indicating that loss of *SCL30a* function does not substantially affect nutrient and water storage in embryonic tissues. Interestingly, the *scl30a-1* mutant also exhibited enhanced seed dormancy. After 7 days at 22 °C in darkness, the germination rate of freshly-harvested, non-stratified *scl30a-1* seeds was only about one third of that of wild-type seeds (Fig 2B). The germination rate of stratified *scl30a-1* mutant seeds was slightly lower, exhibiting a significant delay when compared to wild-type seeds (Fig 2C).

**Fig 2.**
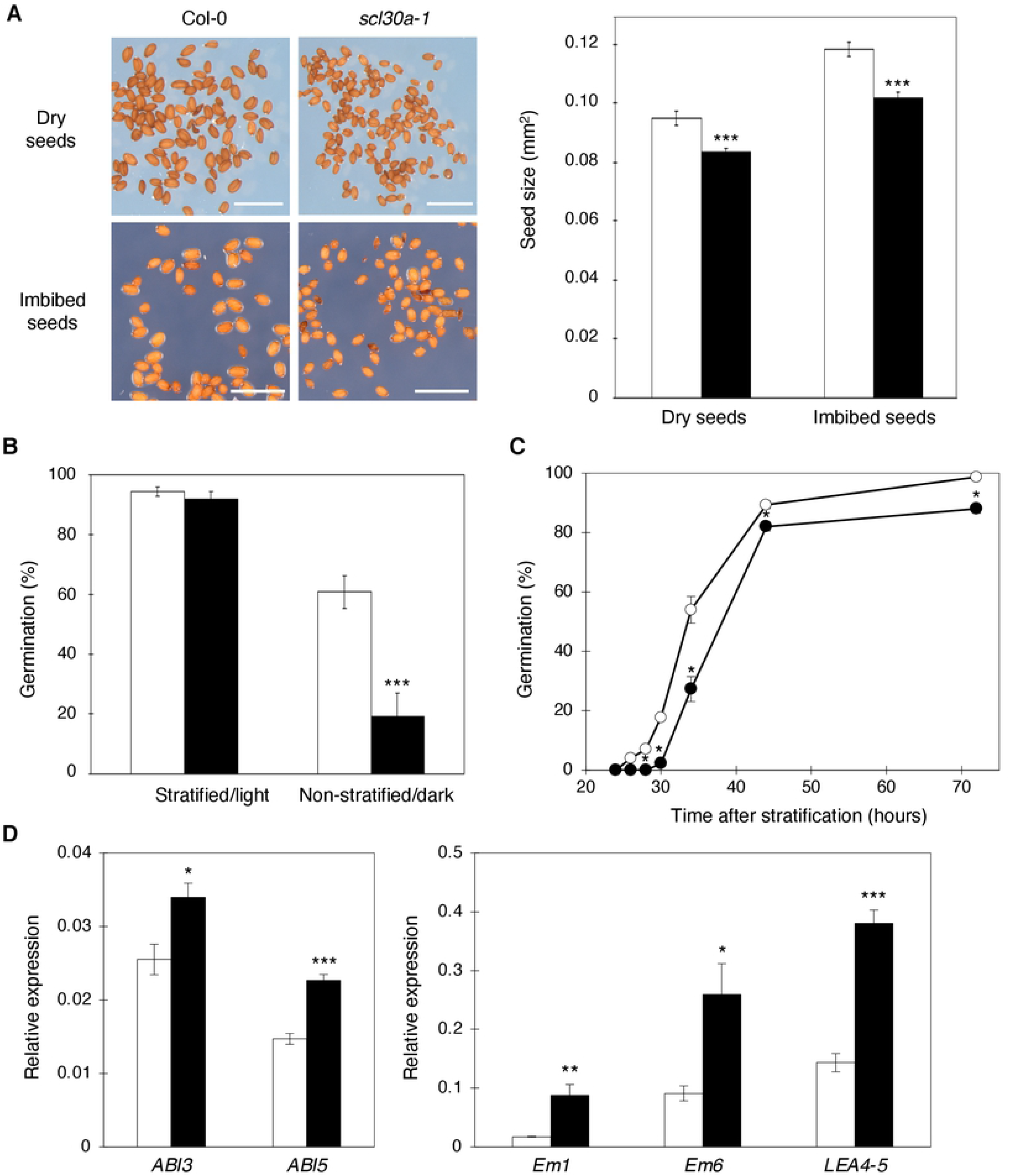
Effect of the *scl30a-1* mutation on seed size, dormancy, germination and seed development gene expression. **(A)** Representative images of dry and imbibed wild-type (Col-0) and mutant (*scl30a-1*) seeds (scale bars, 1.5 mm), and quantification of the area of Col-0 (white bars) and *scl30a-1* (black bars) seeds (means ± SE, *n* ≥ 60). **(B)** Germination percentages of freshly-harvested Col-0 (white bars) and *scl30a-1* (black bars) seeds scored upon either stratification and 7 days of incubation in light (control) or no stratification and 7 days of incubation in darkness (means ± SE, *n* = 3). **(C)** Germination rates of Col-0 (white circles) and *scl30a-1* (black circles) seeds scored during the first 3 days after stratification (means ± SE, *n* = 3). **(D)** RT-qPCR analysis of the expression levels of the *ABI3*, *ABI5*, *Em1*, *Em6* and *LEA4-5* genes in Col-0 (white bars) and *scl30a-1* (black bars) seeds 18 hours after stratification (means ± SE, *n* = 4). Expression of the *cyclophilin* (*ROC5*) gene was used as a loading control. In **A-D**, asterisks indicate significant differences from the Col-0 wild type (* p < 0.05, ** p < 0.01, *** p < 0.001; Student’s *t*-test).

The seed phenotypes of the *scl30a-1* mutant prompted us to analyze the expression of the *ABI3* and *ABI5* genes, two major transcriptional regulators controlling seed development, dormancy and germination [37,38]. RT-qPCR analyses of germinating seeds showed that the expression of *ABI5*, and to a lesser extent also *ABI3*, is significantly increased in the *scl30a-1* mutant (Fig 2D). In agreement, the expression of *Em1*, *Em6*, and *LEA4-5*, three downstream targets of the ABI3 and ABI5 transcription factors [39,40], was also upregulated in *scl30a-1*, even to a larger extent (Fig 2D).

These findings indicate that the SCL30a SR protein plays an *in vivo* role in embryonic tissues, where it affects seed size, dormancy and germination, and controls the expression of key genes regulating seed development and germination.

### The SCL30a protein affects alternative splicing of a small set of genes during seed germination

To gain insight into the molecular functions of the SCL30a RNA-binding protein, we next conducted an RNA-sequencing (RNA-seq) experiment to compare the transcriptomes of wild-type and mutant germinating seeds. We used the Illumina HiSeq 2500 system to sequence mRNA libraries prepared from Col-0 and *scl30a-1* seeds 18 h after stratification and obtained a minimum of 90 million paired-end clean sequence reads per sample. Given the conserved role of SR proteins in pre-mRNA splicing, we first analyzed the alternative splicing changes caused by the *scl30a-1* mutation.

At the time point sampled, we found only 22 alternative splicing events in 21 genes to be differentially regulated in *scl30a-1* mutant seeds (ΔPSI > |15|): seven intron retention (IR), three exon skipping (ES) and 12 alternative 3’(Alt3) or alternative 5’ (Alt5) splice site events (Table 1). Although of low magnitude (ΔPSI < 25 in all cases), the RNA-seq alternative splicing changes were confirmed in the four events selected for validation by RT-PCR, using wild-type and *scl30a-1* RNA samples independent from those analyzed by RNA-seq (S2 Fig). Interestingly, all of the seven differentially-regulated IR events showed lower inclusion levels in the *scl30a-1* mutant, suggesting that SCL30a negatively regulates splicing of these introns. On the other hand, for the three differential ES events identified, the exons were more included in the wild type, pointing to a role of SCL30a in promoting splice site recognition. Thus, IR and ES results point to a contradictory role of this protein in splicing regulation; however, the number of alternative splicing events retrieved is too low to draw conclusions on the mechanistic function of SCL30a.

**Table 1.** Genes displaying alternative splicing changes in germinating *scl30a-1* mutant seeds. Means of percent spliced-in index (PSI) in Col-0 wild-type (WT) and *scl30a-1* mutant germinating seeds (*n* = 3) are presented. Genes are ordered by type of alternative splicing (AS) event — intron retention, (IR, in red), exon skipping (ES, in orange), alternative 5’ splice site (Alt5, in blue) and alternative 3’ splice site (Alt3, in green) — and decreasing absolute ΔPSI values.

| Gene ID | Gene name | AS event ID | AS event type | WT PSI | *scl30a-1* PSI | $\Delta\text{PSI}$ |
| --- | --- | --- | --- | --- | --- | --- |
| AT3G07420 | *SYNC2* | AthINT0106607 | IR | 54.1 | 35.1 | 19.0 |
| AT2G19910 | *RDR3* | AthINT0098183 | IR | 29.1 | 12.2 | 16.9 |
| AT3G07890 |  | AthINT0022974 | IR | 26.0 | 9.2 | 16.8 |
| AT1G06500 |  | AthINT0001122 | IR | 53.1 | 36.6 | 16.5 |
| AT2G46915 |  | AthINT0021222 | IR | 39.0 | 22.5 | 16.5 |
| AT5G64980 |  | AthINT0051338 | IR | 37.1 | 20.7 | 16.4 |
| AT5G09690 | *MGT7/MRS2-7* | AthINT0086682 | IR | 27.8 | 12.2 | 15.6 |
| AT3G63445 |  | AthEX0008460 | ES | 97.3 | 72.4 | 24.9 |
| AT1G45248 |  | AthEX0002068 | ES | 70.5 | 52.9 | 17.6 |
| AT1G10890 |  | AthEX0000589 | ES | 42.4 | 25.4 | 17.0 |
| AT1G15410 |  | AthALTD0000759-2/2 | Alt5 | 89.0 | 68.6 | 20.3 |
| AT1G34340 |  | AthALTD0001643-2/3 | Alt5 | 78.8 | 94.9 | -16.1 |
| AT5G24735 | *SORF31* | AthALTD0009809-7/10 | Alt5 | 32.8 | 48.8 | -16.0 |
| AT5G24735 | *SORF31* | AthALTD0009809-6/10 | Alt5 | 63.8 | 48.7 | 15.2 |
| AT3G05510 |  | AthALTA0008736-3/4 | Alt3 | 54.8 | 33.9 | 20.9 |
| AT3G53470 |  | AthALTA0011304-2/2 | Alt3 | 34.5 | 55.1 | -20.5 |
| AT3G22260 |  | AthALTA0010043-2/2 | Alt3 | 36.2 | 54.6 | -18.5 |
| AT2G05210 | *POT1A* | AthALTA0038411-3/4 | Alt3 | 24.3 | 42.6 | -18.3 |
| AT1G06590 | *APC5* | AthALTA0022122-3/4 | Alt3 | 56.6 | 72.6 | -16.0 |
| AT3G27460 | *SGF29A* | AthALTA0010423-2/4 | Alt3 | 25.7 | 41.2 | -15.6 |
| AT4G38330 |  | AthALTA0014931-4/4 | Alt3 | 74.7 | 59.2 | 15.5 |
| AT5G47380 |  | AthALTA0018392-3/3 | Alt3 | 42.6 | 27.4 | 15.2 |

Of the 21 genes differentially spliced in the *scl30a-1* mutant, only three have been characterized previously: *MRS2-7* encodes a magnesium transporter [41], *POT1a* a DNA-binding protein required for telomere maintenance [42] known to be regulated by alternative splicing [43], and *SGF29a* is a transcriptional co-activator implicated in salt stress responses [44]. Based on the gene annotation at TAIR (https://www.arabidopsis.org), another four genes appear to be also involved in transcription or different aspects of RNA metabolism, while seven play putative roles in many different processes, including lipid or nitrogen metabolism, glycolysis, cell division and protein deubiquitination. Yet, one third of the genes whose splicing was found to be affected by the SCL30a SR protein (i.e., seven genes) are of unknown function.

### The SCL30a protein regulates transcriptional responses related to seed germination and ABA

We next analyzed the gene expression changes caused by loss of function of the *SCL30a* gene. Our RNA-seq analysis revealed 382 genes whose expression was significantly changed by at least two-fold in the *scl30a-1* mutant. Among these, 315 displayed higher transcript levels than the wild type, whereas 67 were downregulated in the *scl30a-1* mutant (S2 Table).

Given the seed and germination phenotypes of the *scl30a-1* mutant (see Fig 2), we then asked whether the genes whose expression was affected by the SCL30a protein were transcriptionally regulated during the seed germination process. To address this question, we quantified the expression levels of the genes up- and downregulated in the *scl30a-1* mutant using data from an extensive germination time-course RNA-seq experiment performed by Narsai et al. [9] (Fig 3). Remarkably, we found that genes repressed by SCL30a (i.e., upregulated in the mutant) are in general highly expressed in dry seeds and downregulated throughout the germination process (Fig 3A). Conversely, genes whose expression is activated by SCL30a (i.e., downregulated in the mutant) show the opposite trend, being lowly expressed in dry seeds and induced during germination (Fig 3B). This finding coincides with the expression pattern of *SCL30a* (see Fig 1) as well as with the delay in germination exhibited by the *scl30a-1* mutant (see Fig 2C) and points to this SR protein as an important positive regulator of seed germination.

**Fig 3.**
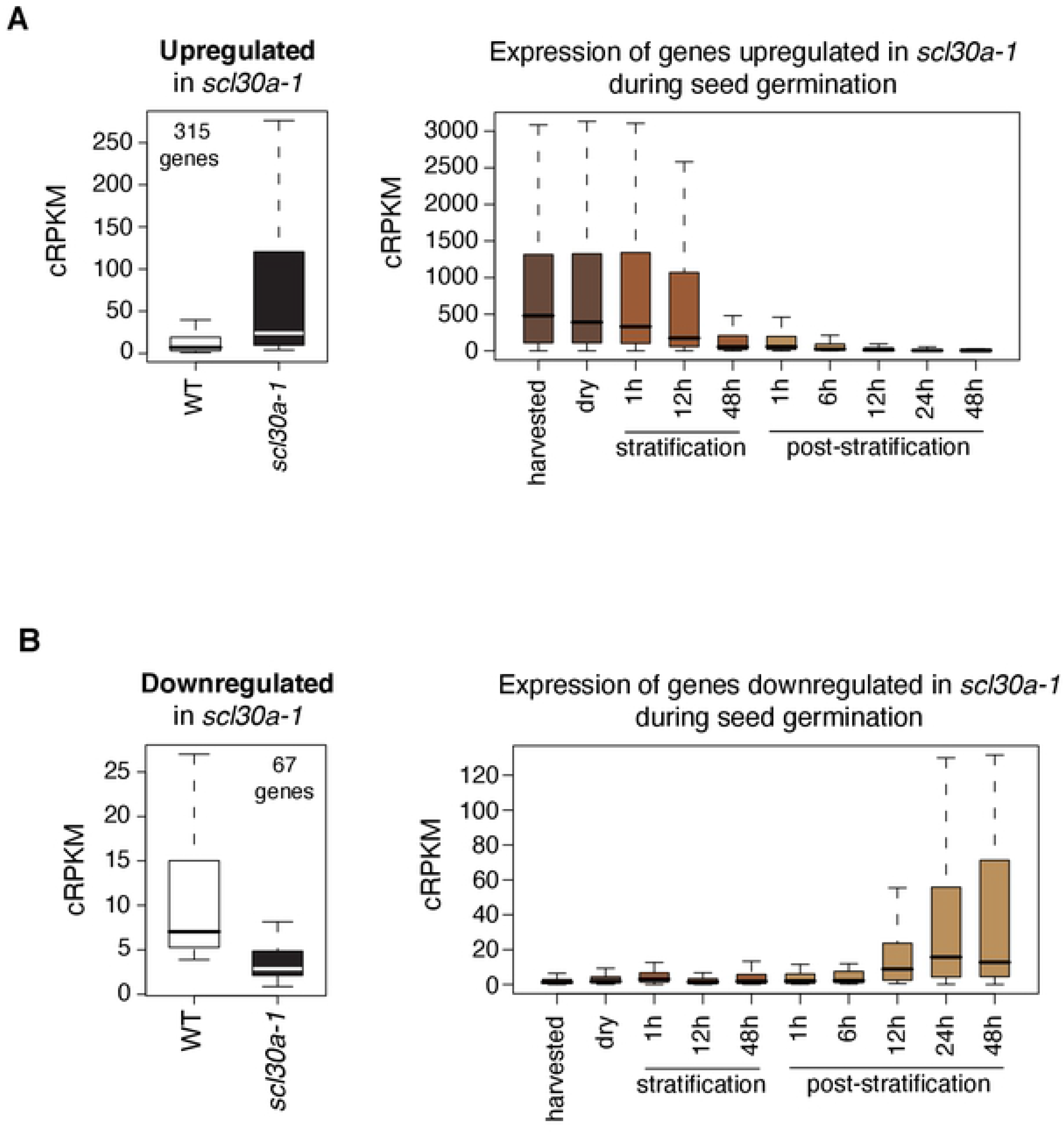
Genes differentially expressed in the *scl30a-1* mutant and their expression pattern during seed germination. Box plot representations of the expression levels of the 315 genes upregulated **(A)** or of the 67 genes downregulated **(B)** in the *scl30a-1* mutant (left) and their expression values in samples collected at different stages of seed germination obtained from [9] (right). See Materials and methods for details.

Importantly, among the genes upregulated in the *scl30a-1* mutant we found many involved in embryo development, seed maturation and dormancy. They include seed storage proteins (e.g., *CRUCIFERIN*) and genes involved in the accumulation and storage of lipidic compounds in seeds (e.g., oleosins), as well as genes involved in the acquisition of desiccation tolerance, such as many *LATE EMBRYOGENESIS ABUNDANT* (*LEA*) genes (S2 Table). In agreement, Gene Ontology (GO) analysis of the *scl30a-1*-upregulated genes showed clear enrichment for categories related to these developmental processes, such as GO:0045735: nutrient reservoir activity, GO:0019915: lipid storage, GO:0009414: response to water deprivation or GO:0009793: embryo development ending in seed dormancy (Fig 4A and S3 Table). Moreover, consistent with the key role played by the ABA hormone in the regulation of seed development, maturation, dormancy and germination, the functional category “GO:0009737: response to abscisic acid” appeared strongly enriched among the *scl30a-1*-upregulated genes. Indeed, the expression of genes encoding main regulators and targets of the ABA signaling pathway — including the ABI5 bZIP transcription factor [45], the seed-specific PP2C AHG1 [46] and the ABA-responsive dehydrin RAB18 [47] — was found to be significantly enhanced in the mutant (Fig 2D and S3 and S4 Tables). On the other hand, many genes found to be downregulated in the *scl30a-1* mutant were related to microtubule activity and cell wall remodeling (Fig 4B and S3 Table), two important processes known to be activated during germination of the seed [1,48].

**Fig 4.**
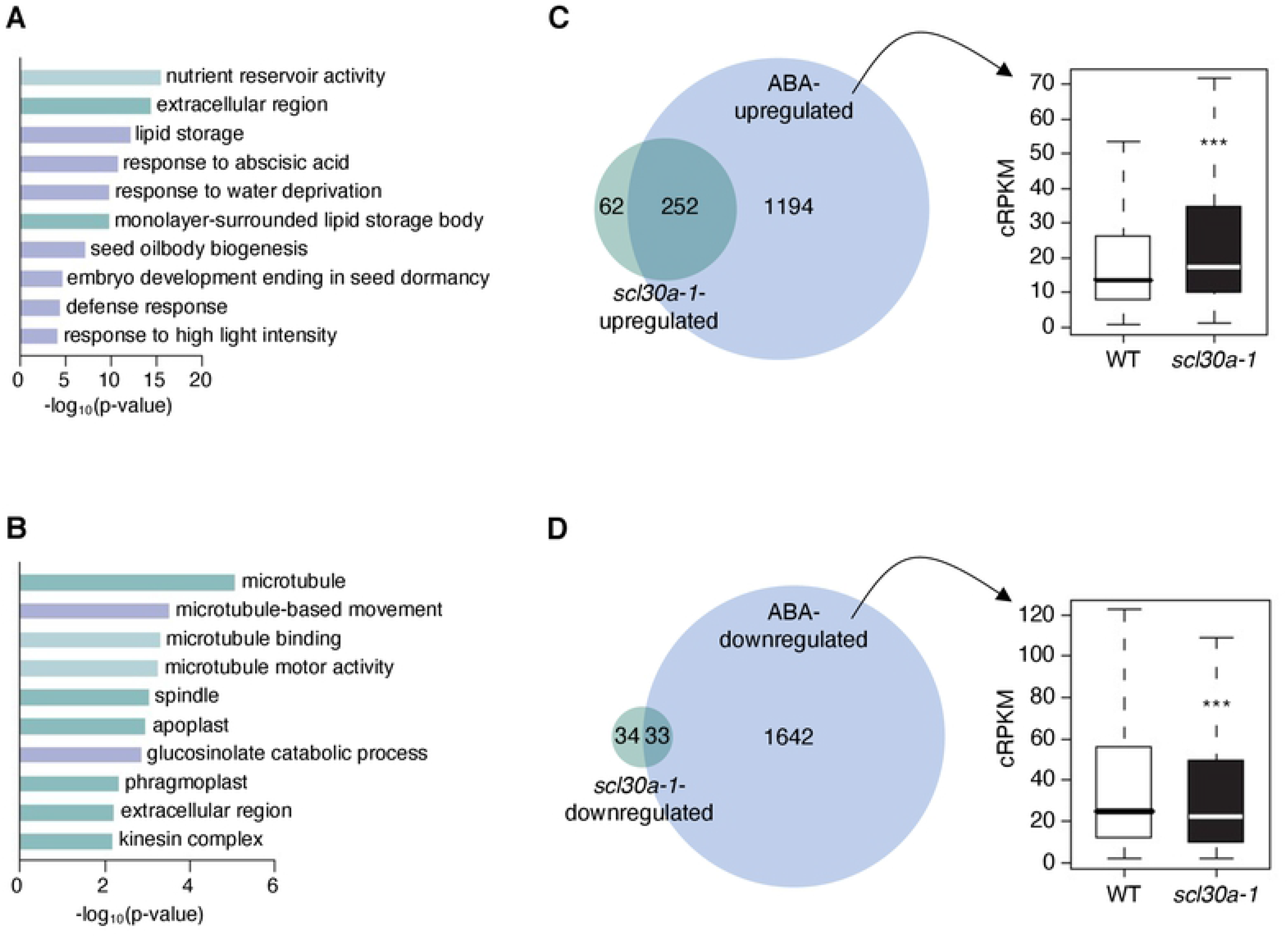
Gene ontology analysis of *scl30a-1*-regulated genes and overlap with ABA transcriptional responses. **(A-B)** The ten most significantly enriched gene ontology categories, including biological process (purple bars), cellular component (dark-green bars) and molecular function (light-green bars), for the genes up- **(A)** and down- **(B)** regulated in the *scl30a-1* mutant. **(C-D)** Overlap between the genes up- **(C)** or down- **(D)** regulated in the *scl30a-1* mutant (green circles) with those up- **(C)** or down- **(D)** regulated by ABA in Arabidopsis germinating seeds from Costa et al. [49] (blue circles), respectively (see Materials and methods for details). The boxplots on the right represent the distribution of expression of the 1446 ABA-upregulated genes **(C)** and 1675 ABA-downregulated genes **(D)** [49] in wild-type (Col-0) and *scl30a-1* mutant germinating seeds (see Materials and methods for details), with the asterisks indicating significant differences from the Col-0 wild type (*Wilcoxon* test, *** p < 0.001).

To gain further insight into the extent of SCL30a control of ABA responses during seed germination, we compared the differentially expressed genes in the *scl30a-1* mutant with a list of ABA-regulated genes obtained from the reanalysis of a previous microarray experiment performed in germinating seeds submitted to a transient ABA treatment [49]. Strikingly, 80 % (252 genes) of the genes upregulated in the *scl30a-1* mutant were also induced by ABA in wild-type germinating seeds (Fig 4C and S5 Table), while 49 % (33 genes) of the genes downregulated in the *scl30a-1* mutant were repressed by ABA (Fig 4D and S6 Table). We then analyzed the expression levels of the ABA-regulated genes defined based on [49] in our RNA-seq data. Interestingly, the 1446 genes upregulated by ABA were significantly more highly expressed in *scl30a-1* than in the wild type, while the 1675 ABA-downregulated genes were downregulated in our mutant (Fig 4C and 4D). Together, these results suggest that an important component of SCL30a function during seed germination is related to the control of ABA-mediated transcriptional responses.

### SCL30a is a positive regulator of ABA signaling and salt stress tolerance during seed germination

To further characterize and confirm the functional role of the SCL30a SR protein in seeds, we generated transgenic Arabidopsis lines expressing the full-length *SCL30a.1* transcript under the control of the 35S promoter in the wild-type Col-0 background. Three independent lines noticeably overexpressing the *SCL30a.1* mRNA, *SCL30a-OX1*, *SCL30a-OX2* and *SCL30a-OX3* (Fig 5A and S3A Fig), were selected for phenotypical characterization. We first assessed the impact of *SCL30a* overexpression on the seed traits found to be affected by the *scl30a-1* mutation (see Fig 2). In contrast to what was observed for the *scl30a-1* mutant, imbibed seeds from the SCL30a-overexpressing plants were significantly (10%) larger than those from wild-type plants (Fig 5B and S3B Fig). Furthermore, stratified *SCL30a*-overexpressing seeds germinated slightly faster under control conditions than wild-type seeds (Fig 5C).

**Fig 5.**
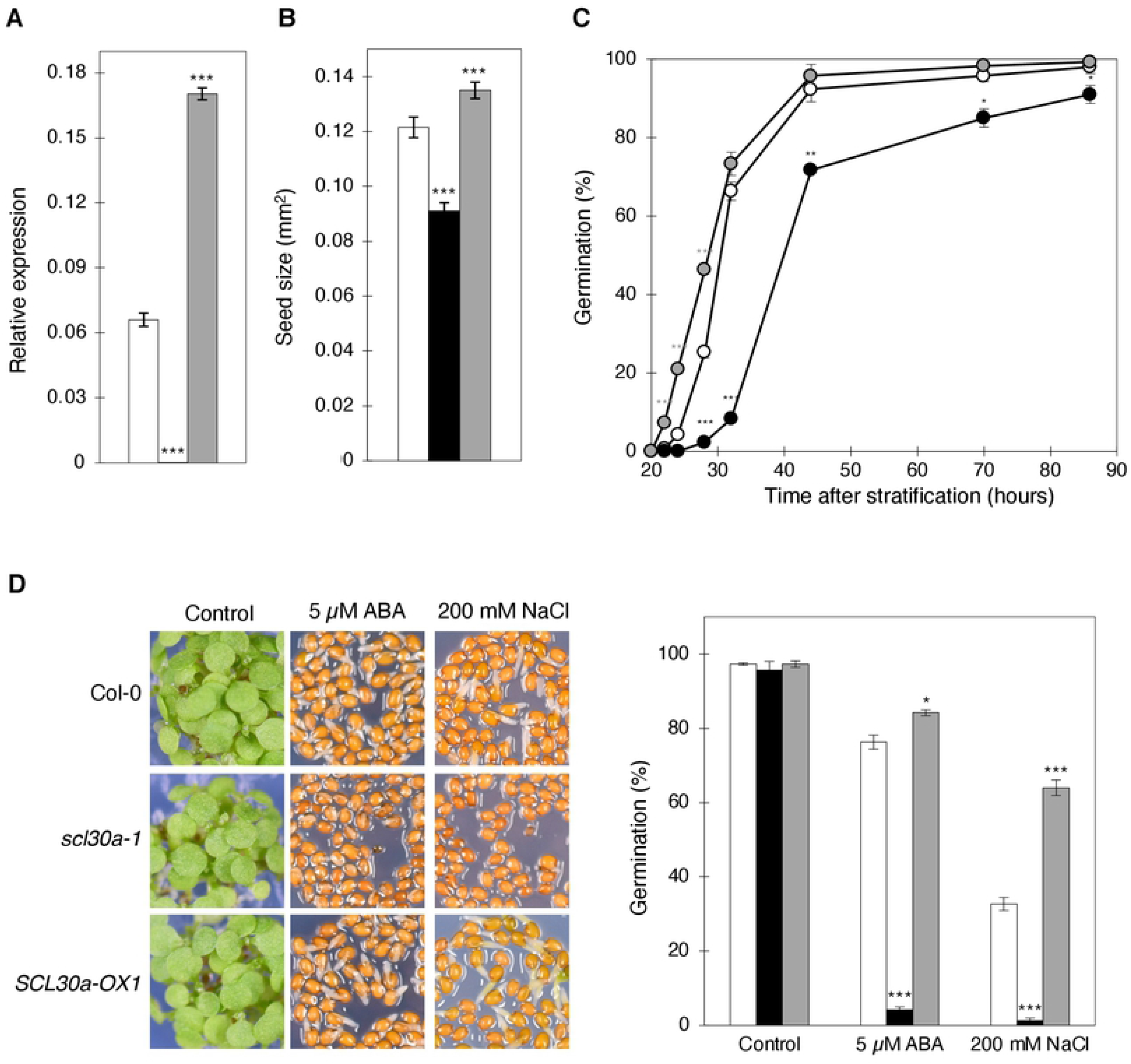
Seed and germination phenotypes conferred by *SCL30a* overexpression. **(A)** RT-qPCR analysis of the expression levels of *SCL30a* in Col-0 (white bar), *scl30a-1* (black bar) and *SCL30a-OX1* (grey bar) 7-day-old seedlings (means ± SE, *n* = 4). Expression of the *ubiquitin* (*UBQ10*) gene was used as a loading control. **(B)** Size (expressed as area) of imbibed Col-0 (white bar), *scl30a-1* (black bar) and *SCL30a-OX1* (gray bar) seeds (means ± SE, *n* ≥ 30). **(C)** Germination rates of Col-0 (white circles), *scl30a-1* (black circles), and *SCL30a-OX1* (gray circles) seeds scored during the first 3.5 days after stratification (means ± SE, *n* = 3). **(D)** Representative images of Col-0, *scl30a-1* and *SCL30a-OX1* seeds germinating in the absence (control) or presence of 5 μM ABA or 200 mM NaCl 7 days after stratification, and germination percentages of Col-0 (white bars), *scl30a-1* (black bars) and *SCL30a-OX1* (gray bars) seeds in the absence (control) or presence of 5 μM ABA or 200 mM NaCl scored 7 days after stratification (means ± SE; *n* = 3). In **A-D**, asterisks indicate statistically significant differences from the Col-0 wild type (* p < 0.05, ** p < 0.01, *** p < 0.001; Student’s *t*-test).

The differential expression of ABA-related genes observed in the *scl30a-1* mutant prompted us to analyze ABA response of the different genotypes during germination. We found that the *scl30a-1* mutant displays strong hypersensitivity to the hormone (Fig 5D and S3C Fig), with less than 10% of the mutant seeds germinating under ABA concentrations that allowed 75% germination of the wild type (Fig 5D). In agreement, seeds from the *SCL30a* overexpression lines were less sensitive to the hormone during seed germination (Fig 5D and S3C Fig). Given the established link between ABA and osmotic stress responses [5], we next examined the effects of loss of function and overexpression of the *SCL30a* gene on seed germination under salt stress. In line with the effect of exogenously applied ABA, the germination rate of mutant seeds in the presence of 200 mM of NaCl was markedly reduced when compared to those of the wild type, while the *SCL30a* overexpression lines were hyposensitive to high salinity, germinating twice as well as the wild type under these conditions (Fig 5D and S3C Fig).

The above findings show that the full-length SCL30a SR protein plays an *in vivo* role in seed development and germination, clearly substantiating the notion that it positively regulates seed size and germination. Moreover, the strikingly opposite phenotypes under ABA and salt stress induced by loss of function and overexpression of SCL30a demonstrate that this Arabidopsis SR protein is a positive regulator of osmotic stress tolerance during germination of the seed.

### SCL30a function in seeds depends on the ABA pathway

To investigate whether the role of SCL30a in salt stress responses is mediated by ABA, we first performed stress germination assays in the presence of fluridone, an inhibitor of ABA biosynthesis [50–52]. Consistent with the well-known role of ABA as a key mediator of salt stress responses [5], addition of 1 μM fluridone notably relieved the inhibition imposed by NaCl on the germination of wild-type seeds (Fig 6A). Most importantly, the presence of fluridone rescued the salt stress hypersensitive phenotype of the *scl30a-1* mutant, which germinated at rates similar to the wild type in NaCl (Fig 6A). This result indicates that the mutant’s salt stress germination phenotype depends on endogenous ABA production.

**Fig 6.**
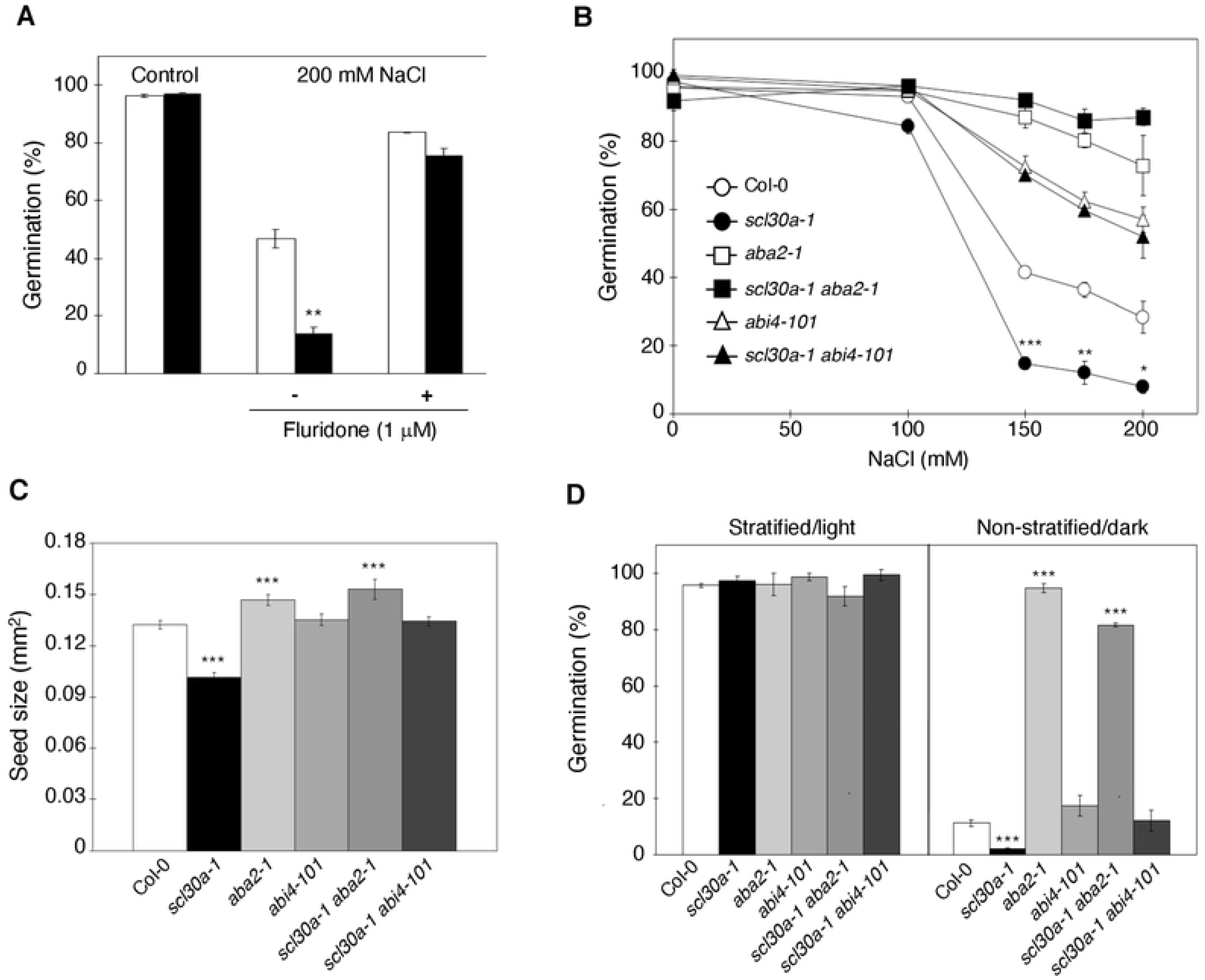
ABA dependence of the *scl30a-1* mutant phenotypes. **(A)** Germination percentages of Col-0 (white bars) and *scl30a-1* (black bars) seeds in the absence (control) or presence of 200 mM NaCl supplemented or not with 1 μM fluridone, scored 5 days after stratification (means ± SE, *n* = 3). Asterisks indicate statistically significant differences from the Col-0 wild type (** p < 0.01; Student’s *t*-test). **(B)** Germination rates of Col-0, *scl30a-1*, *aba2-1*, *abi4-101, scl30a-1 aba2-1* and *scl30a-1 abi4-101* seeds under different NaCl concentrations scored 4 days after stratification (means ± SE, *n* = 3). Asterisks indicate statistically significant differences between the *scl30a-1* mutant and the Col-0 wild type or the double mutants and the corresponding ABA single mutant (* p < 0.05, ** p < 0.01, *** p < 0.001; Student’s *t*-test). **(C)** Size (expressed as area) of imbibed Col-0, *scl30a-1*, *aba2-1*, *abi4-101, scl30a-1 aba2-1* and *scl30a-1 aba4-101* seeds (means ± SE, *n* ≥ 60). Asterisks indicate statistically significant differences from the Col-0 wild type (*** p < 0.001; Student’s *t*-test). **(D)** Germination percentages of freshly-harvested Col-0, *scl30a-1*, *aba2-1*, *abi4-101, scl30a-1 aba2-1* and *scl30a-1 abi4-101* seeds scored upon either stratification and 7 days of incubation in light or 7 days of incubation in darkness (means ± SE, *n* = 3). Asterisks indicate statistically significant differences from the Col-0 wild type (*** p < 0.001; Student’s *t*-test).

To conclusively establish the ABA dependence of SCL30a function, we next turned to epistatic analyses and assessed the genetic interaction between *SCL30a* and *ABA2*, encoding a cytosolic short-chain dehydrogenase reductase involved in the conversion of xanthoxin to ABA-aldehyde during ABA biosynthesis [53], or *ABI4*, which encodes an ERF/AP2-type transcription factor involved in ABA signal transduction [54,55]. To this end, the *scl30a-1* mutant was independently crossed with the ABA-deficient *aba2-1* [6] and ABA-insensitive *abi4-101* [56] mutant alleles to generate the corresponding homozygous double mutants. As seen in the dose-response curves depicted in Fig 6B, seeds from the *scl30a-1aba2-1* and *scl30a-1abi4-101* double mutants behaved as those of the corresponding single ABA mutants when germinated under high salinity, showing that SCL30a control of this stress response fully relies on functional *ABA2* and *ABI4* genes.

We then assessed the seed size and dormancy of the different genotypes. Both the *aba2-1* and the *abi4-101* mutations suppressed the reduced size displayed by *scl30a-1* seeds, with the area of *scl30a-1aba2-1* imbibed seeds being even significantly larger than those of the wild type, as previously reported for the *aba2-1* mutant [8] (Fig 6C). Regarding seed dormancy, the double mutants again showed strikingly similar phenotypes to those induced by single mutations in the *ABA2* and *ABI4* genes that, in agreement with early reports [6,57], conferred strongly reduced and normal dormancy, respectively (Fig 6D). Therefore, both *ABA2* and *ABI4* are epistatic to the *SCL30a* gene, indicating that the seed/germination roles of the encoded SR protein are fully dependent on a functional ABA pathway.

The above findings raised the question of whether changes in SCL30a levels affect ABA biosynthesis or sensing/signaling of the stress hormone. To address this issue, we measured the endogenous ABA content of wild-type, *scl30a-1* mutant and *SCL30a*-overexpressing seeds germinated in control conditions or under high salinity stress. Table 2 shows that Col-0, *scl30a-1* and *SCL30a-OX* seeds responded to the presence of 200 mM NaCl by increasing their ABA content by around two-fold, with no significant differences in ABA levels being observed between the three genotypes either in the absence or presence of salt stress. As expected, the ABA content of the ABA biosynthesis *aba2-1* mutant, included as a negative control, was unaltered by high salinity stress (Table 2). These results suggest that SCL30a activity does not influence endogenous ABA levels in seeds, rather affecting sensing and/or signal transduction of the hormone during seed germination.

**Table 2.** Effect of loss of function or overexpression of *SCL30a* on seed ABA levels. ABA content (means ± SE, *n* = 6-8), in ng/g of fresh weight, of Col-0, *scl30a-1*, *SCL30a-OX2* and *aba2-1* seeds germinated for 2 days in the absence or presence of 200 mM NaCl. Letters indicate significantly different ABA levels between genotypes among each condition and asterisks significant differences for each genotype between control and salt stress conditions (p < 0.05; Student’s *t*-test).

| Genotype | Control | NaCl | NaCl/Control |
| --- | --- | --- | --- |
| Col-0 | 31.62 ± 3.07 | 59.06 ± 8.40 | *1.86 ± 0.40 |
| *scl30a-1* | 36.45 ± 7.40 | 73.04 ± 16.60 | *2.00 ± 0.80 |
| *SCL30a-OX2* | 36.54 ± 7.29 | 89.00 ± 19.91 | *2.44 ± 0.98 |
| *aba2-1* | 27.86 ± 6.61 | 24.25 ± 4.11<sup>a</sup> | 0.87 ± 0.35 |

## Discussion

The first indication that the Arabidopsis SCL30a SR protein was involved in regulating seed-specific traits came from our gene expression studies, showing high *SCL30a* induction in the embryo and testa of imbibed seeds as well as during the first stages of germination. Phenotypical characterization of the *scl30a-1* loss-of-function mutant then revealed that this gene is required to achieve the final size and adequate dormancy levels of mature Arabidopsis seeds, as well as subsequently during the germination process. Importantly, we also show that SCL30a negatively regulates the response to salt stress as well as ABA signaling during germination of the seed. Accordingly, germinating *scl30a-1* mutant seeds display higher expression of ABA-related genes, and overexpression of *SCL30a* results in a drastic reduction of seed sensitivity to high salinity, corroborating a role for this protein as a positive regulator of abiotic stress tolerance during seed germination.

Although the *SCL30a* gene displays ubiquitous expression in vegetative tissues, we were unable to identify any evident phenotype at later developmental stages. This is likely due to functional redundancy between members of the SCL subfamily at the adult stage. In fact, previous phenotypic studies of adult Arabidopsis plants from single mutants in individual SCL genes did not report any visible alterations, with only a quintuple mutant of the four SCL members and the *SC35* gene (*scl28 scl30 scl30a scl33 sc35*) exhibiting clear defects in leaf development and flowering [33].

Physiological assays using an ABA biosynthesis inhibitor and epistatic analyses with the ABA-biosynthesis *ABA2* [6] and the ABA-signaling *ABI4* [57] genes demonstrate that SCL30a regulation of seed traits is fully dependent on an intact ABA pathway. This is consistent with the global transcriptional changes associated with the loss of *SCL30a* function, showing a clear enrichment of ABA-related functions among the genes upregulated in the *scl30a-1* mutant. Moreover, unchanged ABA levels in mutant and overexpressing seeds, together with the enhanced and reduced sensitivity to exogenously applied ABA caused respectively by loss-of-function and overexpression of *SCL30a*, indicate that the encoded SR protein represses signal transduction of the phytohormone rather than its biosynthesis.

While the central roles of ABA in the induction and maintenance of seed dormancy as well as in mediating responses to salt stress are well established [5], few studies have addressed the involvement of this phytohormone in determining seed size. Nonetheless, expression of the ubiquitin interaction motif-containing DA1 protein, which limits seed size by restricting the period of cell proliferation in the seed integuments, is induced by ABA and a *da1* mutant allele displays altered ABA sensitivity. However, unlike SCL30a, DA1 function appears to be independent of the *ABI4* gene [58]. ABA has also been reported to regulate final seed size via the control of endosperm cellularization during seed development, as reflected by the larger seeds of the *aba2* and *abi5* mutants [8]. Given the smaller seeds produced by the *scl30a-1* mutant and the newly discovered role for SCL30a as a major regulator of ABA transcriptional responses, it appears more likely that this SR protein regulates endosperm development, and thereby seed size, by controlling the expression of key ABA components such as the *ABI5* gene, which is upregulated in the *scl30a-1* mutant.

Seeds challenged with osmotic stress undergo an arrest in germination that is triggered by a rise in their ABA content [59,60]. Our results indicate that by decreasing sensitivity to this phytohormone, the SCL30a SR protein enhances salt stress tolerance during seed germination. The derepression of a subset of ABA-response genes and the germination delay associated with the loss of SCL30a function in the absence of stress suggest that the SR protein is already able to repress ABA signaling under optimal growth conditions. Therefore, it is possible that the hypersensitive phenotype of the *scl30a-1* mutant is a consequence of an already active ABA signaling state, with the stress stimulus inducing an overaccumulation of ABA-responsive transcripts in the mutant. Alternatively, the stronger phenotype of *scl30a-1* under stress when compared to control conditions could indicate stress regulation of SCL30a activity. The fact that the *SCL30a* expression and splicing pattern is unaffected by ABA or salt (data not shown and [34,49]) points to posttranslational regulation of this RNA-binding protein. In support of this notion, SR proteins are known to undergo extensive phosphorylation at their RS domain [14], and stress cues affect both the phosphorylation status and activity of Arabidopsis SR and SR-related proteins [23,61–65]. Notably, SR protein kinase 4 (SRPK4) and stress-responsive mitogen-activated protein kinases (MAPKs) were found to phosphorylate SCL30, a close SCL30a paralog [66].

Quite surprisingly, our large-scale transcriptome analysis revealed only 22 alternative splicing events in 21 genes affected in the *scl30a-1* mutant (dPSI > |15|), thus precluding solid mechanistic insight into the splicing function of this SR protein. Our results contrast with a main expected role for SCL30a as a splicing regulator and raise the question of whether this protein is involved in regulating other steps of gene expression. Beyond splicing, animal SR proteins have been shown to play important roles in coordinating several steps of gene expression, including transcriptional activation, nonsense-mediated decay, mRNA export and translation [18,19,67]. In Arabidopsis, SCL proteins can interact with the NRPB4 subunit of the RNA Polymerase II, pointing to a potential role in the regulation of gene transcription, and simultaneous disruption of the four SCL subfamily genes and *SC35* causes drastic transcriptional changes [33]. Therefore, and in alignment with our transcriptomic results, an important component of SCL30a function during seed germination could lie in the regulation of gene transcription. Nonetheless, the RNA-seq experiment performed here reflects the transcriptome of *scl30a-1* germinating seeds at a specific time point (18 hours after stratification), and the possibility that the observed gene expression changes are a consequence of an earlier alternative splicing defect cannot be ruled out. Future identification of the direct targets of SCL30a using immunoprecipitation methods should provide insight into the molecular functions of this protein during seed germination and stress responses.

Seed size is a major component of crop yield and salt stress dramatically reduces plant productivity worldwide. We have disclosed a novel function for an Arabidopsis splicing factor —SCL30a — in governing seed size and tolerance to salt stress. Our data also suggest a non-canonical role for the SCL30a protein, where it could regulate gene transcription rather than alternative splicing during seed germination. Moreover, we provide evidence that SCL30a modulates seed traits by interacting with the ABA pathway. The larger and salt-tolerant seeds produced by SCL30a-overexpressing plants underscore the high potential of this protein for biotechnical applications. Deeper insight into the mode of action of SCL30a may translate into improved crop performance under adverse environmental conditions.

## Materials and methods

### Plant materials and growth conditions

The *Arabidopsis thaliana* ecotype Colombia (Col-0) was used as the wild type in all experiments. Seeds were surface-sterilized for 10 minutes in 50 % (v/v) bleach and 0.07% (v/v) TWEEN^®^20, stratified for 3 days at 4 °C in the dark and plated on MS media [1X Murashige and Skoog (MS) salts (Duchefa Biochemie), 2.5 mM MES (pH 5.7), 0.5 mM myo-inositol and 0.8 % (w/v) agar], before transfer to a growth chamber under 16-h photoperiod (long-day conditions) or continuous light (cool white fluorescent bulbs, 18W840, 4000K at 100 μmol m^−2^ s^−1^) at 22 °C (light period) or 18 °C (dark period) and 60 % relative humidity. Seed imbibition (Fig 1, 2A, 5B and 6C) was always performed at 4 °C (equivalent to stratification). After 2-3 weeks, seedlings were transferred to soil in individual pots.

PCR-based genotyping of the SALK_041849 line (obtained from NASC) with primers specific for *SCL30a* and the left border of the T-DNA (S7 Table) followed by sequencing of the genomic DNA/T-DNA junction confirmed the insertion site and allowed isolation of a homozygous line, which was backcrossed twice with the wild type. The *scl30a-1* mutant was independently crossed with the *aba2-1* [6] and the *abi4-101* [57] alleles (obtained from NASC) and the corresponding double mutants identified via PCR screening (S7 Table) of F2 progeny following F1 self-fertilization.

### Generation of transgenic plants

Plant transformation was achieved by the floral dip method [68] using *Agrobacterium tumefaciens* strain EHA105.

For reporter gene experiments, the 2206 bp immediately upstream of the *SCL30a* start codon were PCR-amplified (S7 Table) from genomic DNA and subcloned into the pGEM vector (Promega), where the eGFP-GUS segment isolated from the pKGWFS7 vector [69] using the *Sac*II/*Nco*I restriction sites was fused at the 3’ end of the *SCL30a* promoter sequence. The entire fragment was transferred into pKGWFS7 via the *Spe*I/*Nco*I restriction sites, replacing the original CmR-ccdB-eGFP-GUS cassette, and the construct agroinfiltrated into Col-0 plants.

To generate the Pro*35S:SCL30a.1* construct, an RT-PCR fragment corresponding to the *SCL30a.1* transcript (S7 Table) was inserted into the pBA002 backbone using the *Asc*I/*Pac*I restriction sites, and the construct agroinfiltrated in Col-0 plants. Two independent *SCL30a-OX* lines were first isolated and analyzed. After several seed-to-seed cycles, expression of the transgene in these *SCL30a-OX2* and *SCL30a-OX3* lines was silenced, with consequent loss of the corresponding phenotypes. A third overexpression line, *SCL30a-OX1*, was then generated and phenotypically characterized.

### Seed measurements and composition

Wild-type (Col-0) and mutant (*scl30a-1*) plants were sown and grown to maturity simultaneously under identical conditions, and all assays were performed with seeds from comparable lots.

The area of dry and imbibed seeds was measured using the ImageJ software (http://rsbweb.nih.gov/ij). To determine seed weight, six groups of 1000 dry seeds were weighed using an Acculab ALC-80.4 (Sartorius) analytical balance.

For compositional analysis, dry seeds were bulk harvested by genotype and homogenized. The oil, protein and soluble carbohydrate contents were determined as described previously [70]. To analyze fatty acids, dry seeds (20 mg) were crushed and sonicated in 2 mL of heptane for 15 minutes at 60 °C. After centrifuging for 5 minutes at 2,000 *g*, 200 μL of the heptane layer were transferred to a small vial with 50 μL of trimethylsulfonium hydroxide (TMSH) in methanol and an additional 300 μL of heptane. After incubation for 30 minutes at room temperature, 1 μL of the upper heptane layer was used to analyze the fatty acid methyl esters, which were separated and quantified using a Hewlett-Packard 6890 gas chromatograph as described in Cahoon *et al.* (2001). All analyses were performed in duplicate on three independent seed batches per genotype.

### Germination and dormancy assays

For germination assays, fully mature siliques from dehydrated plants were collected and stored in the dark at room temperature for at least one week before phenotypical analysis. After surface-sterilization and stratification for 3 days at 4 °C in the dark, 70-100 seeds of each genotype were sown on MS media supplemented or not with the appropriate concentrations of NaCl, ABA (mixed isomers, A1049; Sigma) or fluridone (45511, Fluka) and then transferred to long-day conditions, except for the determination of germination rates under control conditions (Fig 2C and 5C), which was conducted under continuous light to avoid the effect of long dark periods during a short time course. To assess dormancy, seeds from freshly mature siliques were collected from the tree, immediately surface-sterilized and plated on MS media before transfer to dark at 22 °C, with control seeds being stratified for 3 days at 4 °C in the dark before transfer to long-day conditions. Percentages of seed germination, defined as protrusion of the radicle through the seed coat, were scored over the total number of seeds. The results presented are representative of at least three independent experiments.

### ABA content determination

Mature seeds harvested from Col-0, *scl30a-1*, *SCL30a-OX2* or *aba2-1* dehydrated plants and stored for 5 months were stratified for 3 days at 4 °C in the dark, sown on MS media with or without 200 mM NaCl and grown for 2 days under long-day conditions. Seeds were then collected and endogenous ABA levels quantified using an immunoassay as described in [26].

### Expression and alternative splicing analyses of individual genes

Histochemical staining of GUS activity in Pro*SCL30a:GUS* transgenic lines was performed as described by Sundaresan *et al.* [71].

For the RT-PCR analysis shown in Fig 1, S1 Fig and S3 Fig, total RNA was extracted from different plant tissues using TRI Reagent (T924; Sigma-Aldrich) or from dry, imbibed and up to 5-day germinated seeds using the innuPREP Plant RNA Kit (Analytik Jena). First-strand cDNA synthesis and PCR amplification were performed as described in [72], using the primers and number of cycles indicated in Table S7 as well as *ROC1* as a reference gene. Results are representative of at least three experiments.

For the RT-qPCR analyses shown in Fig 2D, seeds were stratified for 3 days at 4 °C, sown on MS media, transferred to continuous light conditions, and collected after 18 hours (prior to radicle emergence) to avoid major developmental effects. For the RT-qPCR of Fig 5A, seedlings were grown for 1 week after stratification. Total RNA was extracted using the innuPREP Plant RNA Kit (Analytik Jena), digested with the RQ1 DNase (Promega), and first strand cDNA synthesized using 1 μg RNA, Super Script III Reverse Transcriptase (Invitrogen) and a poly-T primer. qPCR was performed using an ABI QuantStudio sequence detection system (Applied Biosystems) and Luminaris Color HiGreen qPCR Master Mix (Thermo Scientific) on 2.5 μL of cDNA (diluted 1:10) per 10-μL reaction volume, containing 300 nM of each gene-specific primer (S7 Table). Reaction cycles were 95 °C for 2 min (1X), 95° C for 30 s/60 °C for 30 s/72 °C for 30 s (40X), followed by a melting curve step to confirm the specificity of the amplified products. *UBQ10* and *ROC5* were used as reference genes. Each experiment was replicated at least three times.

For the analyses of alternative splicing shown in S2 Fig, PCR with the NZYTaq II 2x Green Master Mix (Nzytech) was performed on cDNA from three biological replicates of germinating seeds (18 hours after stratification, continuous light) using primers flanking the alternatively spliced intron (S7 Table) obtained from PASTDB (pastdb.crg.edu). Reaction cycles were 95 °C for 3 min (1X), 95 °C for 30 s/58 °C for 30 s/72 °C for 5 min (35X). The PCRs products then were loaded on a 2% agarose gel and gel bands quantified using the ImageJ software (http://rsbweb.nih.gov/ij). The percent spliced-in (PSI) for each alternative splicing event was calculated after quantification of the inclusion (I) or splicing (S) for a given event as PSI = I / (I + S).

### RNA-seq sample preparation and sequencing

Approximately 50 mg of Col-0 wild type and *scl30a-1* mutant seeds (three biological replicates) were surface-sterilized, stratified at 4 °C for 3 days and sown on MS media for 18 hours under continuous light, before total RNA was extracted using the innuPREP Plant RNA Kit (Analytik Jena). The RNA-seq libraries generated from Col-0 and *scl30a-1* seeds were prepared and sequenced at the Center for Genomic Regulation (Barcelona, Spain) using the HiSeq Sequencing V4 Chemistry kit (Illumina, Inc) and the HiSeq 2500 sequencer (Illumina, Inc), with a read length of 2 × 125 bp.

### RNA-seq quantification of sequence inclusion and identification of differentially-spliced genes

We employed vast-tools v2.5.1 to quantify alternative splicing from RNA-seq for *A. thaliana* [36,73]. This tool quantifies exon skipping (ES), intron retention (IR), and alternative 3’ (Alt3) and 5’ (Alt5) splice sites. For all these types of events, vast-tools estimates the percent of inclusion of the alternative sequence (PSI) using only exon-exon (or exon-intron for IR) junction reads and provides information about the read coverage (see https://github.com/vastgroup/vast-tools for details). To identify alternative splicing events regulated by SCL30a we used vast-tools compare. This function compared PSI values of each AS event with sufficient read coverage in all wild-type and *scl30a-1* samples being tested (three biological replicates of each genotype) and selected those with an average ΔPSI > 15 and a ΔPSI between the two distributions > 5 (--min_dPSI 15 --min_range 5) (see https://github.com/vastgroup/vast-tools and [73] for details). We also used the --p_IR filter to discard introns with a significant read imbalance between the two exon-intron juntions (p < 0.05, binomial test; see [74] for details). Moreover, to ensure that Alt3 and Alt5 are located in exons with a sufficient inclusion level, we used the option –min_ALT_use 25, which implies that the host exon has a minimum PSI of at least 25 in each analyzed sample.

### RNA-seq quantification of gene expression and identification of differentially expressed genes

Quantification of Arabidopsis transcript expression from our RNA-seq experiment and public sequencing data on seed germination (GSE94459) was performed using vast-tools v2.5.1 [73]. This tool provides cRPKMs numbers for each Arabidopsis transcript as the number of mapped reads per million mapped reads divided by the number of uniquely mappable positions of the transcript [36]. To identify differentially expressed genes between wild-type and *scl30a-1* germinating seeds, we used vast-tools compare_expr using the option -norm (see https://github.com/vastgroup/vast-tools for details). In brief, a quantile normalization of cRPKM values with “Normalize Between Arrays” within the “limma” package of R is first performed. Next, genes that were not expressed at cRPKM > 5 are filtered out and read counts > 50 across all the replicates of at least one of the genotypes compared. Graphs in Fig 3 and 4 only show expression of genes that passed these cut-offs. Finally, differentially-expressed genes were defined as those with a fold change of at least 2 between each of the individual replicates from each genotype.

### Assessment of overlap between SCL30a- and ABA-regulated genes

ABA-regulated genes were obtained from the reanalysis of GSE62876 [49] using the default settings of GEO2R (https://www.ncbi.nlm.nih.gov/geo/info/geo2r.html). Three comparisons were conducted: 0 hours of ABA treatment versus 2, 12 or 24 hours. Genes regulated at any of these timepoints were selected (FC > 2; adjusted p-value < 0.05). Given that ABA-regulated genes in [49] are defined based on microarray studies, which do not assess expression of all Arabidopsis genes as RNA-seq experiments do, for the comparison we discarded one SCL30a-regulated gene not represented in the microarray. We also discarded ABA-regulated genes not expressed in our RNA-seq samples (see previous section).

### Gene ontology enrichment analyses

The Gene Ontology (GO) enrichment analysis shown in Fig 4, which identifies significantly enriched biological processes, molecular functions and cellular components among the genes up- and downregulated in the *scl30-1* mutant, was performed using the functional annotation classification system DAVID version 6.8 [75]. Only statistically significant GO categories (p < 0.05) are shown in Table S3.

### Accession numbers

Raw sequencing data and transcript expression results were submitted to the Sequence Read Archive (accession number GSE181122).

## Acknowledgments

We thank V. Nunes (IGC Plant Facility) for technical assistance and excellent plant care. This work was funded by Fundação para a Ciência e a Tecnologia (FCT) through Grants PTDC/AGR-PRO/119058/2010 and BIA-FBT/31018/2017, as well as PhD Fellowship SFRH/BD/28519/2006 awarded to S.D.C. Funding from the research unit GREEN-it “Bioresources for Sustainability” (UIDB/04551/2020) is also acknowledged. T.L was supported by Marie Skłodowska-Curie Individual Fellowship MSCA-IF-2015 (project 706274), and G.M. was supported by MSCA-IF-2016 (project 750469) and an EMBO Long-Term Fellowship (ALTF 1576-2016).

## Supporting Information Legends

**S1 Fig. Structure of the *SCL30a* gene and isolation of the *scl30a-1* loss-of-function mutant.**

**(A)** Schematic representation of the *SCL30a* gene showing the site of insertion and orientation of the T-DNA in the *scl30a-1* mutant (boxes indicate exons with UTRs in grey, lines between boxes represent introns, and arrows indicate the location of *SCL30a*- and T-DNA-specific primers), and structure of the three identified splice variants as well as of the corresponding predicted protein isoforms (RRM, RNA recognition motif; RS, arginine/serine-rich domain). The asterisks mark the position of the predicted protein truncation in the *scl30a-1* mutant. **(B)** RT-PCR analysis of *SCL30a* transcript levels in wild-type (Col-0) and mutant (*scl30a-1)* 5-day old seedlings using primers flanking the T-DNA, and up- or downstream of the insertion site. The location of the F1, F3, R1 and R2 primers is shown in (A). The *UBIQUITIN 10* (*UBQ10*) gene was used as a loading control.

**S2 Fig. Validation of selected differential alternative splicing events detected by RNA-seq.**

RT-PCR analysis of individual AS events differentially regulated between Col-0 and *scl30a-1* germinating seeds 18 hours after stratification in **(A)** AT5G64980 (event: AthINT0051338), **(B)** AT3G07890 (event: AthINT0022974), **(C)** AT2G46915 (event: AthINT0021222) and **(D)** AT5G09690 (event: AthINT0086682). Graphs represent percent spliced-in (PSI) values (means ± SE *n* = 3-5) after quantification of the corresponding band intensities using the Image J software. Asterisks indicate statistically significant differences from the Col-0 wild type (* p < 0.05; Student’s *t*-test).

**S3 Fig. Characterization of the *SCL30a-OX2* and *SCL30a-OX3* overexpression lines.**

**(A)** RT-PCR analysis of *SCL30a* transcript levels in 7-day-old seedlings of Col-0, *scl30a-1* and two *SCL30a* overexpression lines (*SCL30a-OX2* and *SCL30a-OX3*). The location of the F2 and R1 primers is shown in S1A Fig. The *UBIQUITIN 10* (*UBQ10*) gene was used as a loading control. **(B)** Size (expressed as area) of imbibed Col-0 (white bars), *scl30a-1* (black bars) and *SCL30a-OX2* or *SCL30a-OX3* (gray bars) seeds (means ± SE, *n* ≥ 60). **(C)** Germination percentages of Col-0 (white bars), *scl30a-1* (black bars) and *SCL30a-OX2* or *SCL30a-OX3* (gray bars) in the absence (control) or presence of 3 μM ABA or 200 mM NaCl scored 5 days after stratification. Bars represent means ± SE, *n* = 3. In **B** and **C**, asterisks indicate significant differences from the Col-0 wild type (* p < 0.05, ** p < 0.01, *** p < 0.001; Student’s *t*-test).

## References

1. Holdsworth MJ, Bentsink L, Soppe WJJ. Molecular networks regulating Arabidopsis seed maturation, after-ripening, dormancy and germination. New Phytologist. 2008;179: 33–54. doi:10.1111/j.1469-8137.2008.02437.x

2. Holdsworth MJ, Finch-Savage WE, Grappin P, Job D. Post-genomics dissection of seed dormancy and germination. Trends in Plant Science. 2008;13: 7–13. doi:10.1016/j.tplants.2007.11.002

3. Shu K, Liu X, Xie Q, He Z. Two Faces of One Seed: Hormonal Regulation of Dormancy and Germination. Molecular Plant. 2016;9: 34–45. doi:10.1016/j.molp.2015.08.010

4. Penfield S. Seed dormancy and germination. Current Biology. 2017;27: R874–R878. doi:10.1016/j.cub.2017.05.050

5. Chen K, Li G-J, Bressan RA, Song C-P, Zhu J-K, Zhao Y. Abscisic acid dynamics, signaling, and functions in plants. Journal of Integrative Plant Biology. 2020;62: 25–54. doi:10.1111/jipb.12899

6. Léon-Kloosterziel KM, Gil MA, Ruijs GJ, Jacobsen SE, Olszewski NE, Schwartz SH, et al. Isolation and characterization of abscisic acid-deficient Arabidopsis mutants at two new loci. The Plant Journal. 1996;10: 655–661. doi:10.1046/j.1365-313X.1996.10040655.x

7. Nakashima K, Fujita Y, Kanamori N, Katagiri T, Umezawa T, Kidokoro S, et al. Three Arabidopsis SnRK2 Protein Kinases, SRK2D/SnRK2.2, SRK2E/SnRK2.6/OST1 and SRK2I/SnRK2.3, Involved in ABA Signaling are Essential for the Control of Seed Development and Dormancy. Plant and Cell Physiology. 2009;50: 1345–1363. doi:10.1093/pcp/pcp083

8. Cheng ZJ, Zhao XY, Shao XX, Wang F, Zhou C, Liu YG, et al. Abscisic Acid Regulates Early Seed Development in Arabidopsis by ABI5-Mediated Transcription of SHORT HYPOCOTYL UNDER BLUE1. The Plant Cell. 2014;26: 1053–1068. doi:10.1105/tpc.113.121566

9. Narsai R, Gouil Q, Secco D, Srivastava A, Karpievitch YV, Liew LC, et al. Extensive transcriptomic and epigenomic remodelling occurs during Arabidopsis thaliana germination. Genome Biology. 2017;18: 172. doi:10.1186/s13059-017-1302-3

10. Laloum T, Martín G, Duque P. Alternative Splicing Control of Abiotic Stress Responses. Trends in Plant Science. 2018;23: 140–150. doi:10.1016/j.tplants.2017.09.019

11. Lou L, Ding L, Wang T, Xiang Y. Emerging Roles of RNA-Binding Proteins in Seed Development and Performance. International Journal of Molecular Sciences. 2020;21. doi:10.3390/ijms21186822

12. Chen W, Moore MJ. Spliceosomes. Current Biology. 2015;25: R181–R183. doi:10.1016/j.cub.2014.11.059

13. Meyer K, Koester T, Staiger D. Pre-mRNA Splicing in Plants: In Vivo Functions of RNA-Binding Proteins Implicated in the Splicing Process. Biomolecules. 2015;5: 1717–1740. doi:10.3390/biom5031717

14. Morton M, AlTamimi N, Butt H, Reddy ASN, Mahfouz M. Serine/Arginine-rich protein family of splicing regulators: New approaches to study splice isoform functions. Plant Science. 2019;283: 127–134. doi:10.1016/j.plantsci.2019.02.017

15. Shepard PJ, Hertel KJ. The SR protein family. Genome Biology. 2009;10: 242. doi:10.1186/gb-2009-10-10-242

16. Zhou Z, Fu X-D. Regulation of splicing by SR proteins and SR protein-specific kinases. Chromosoma. 2013;122: 191–207. doi:10.1007/s00412-013-0407-z

17. Barta A, Kalyna M, Lorković ZJ. Plant SR Proteins and Their Functions. In: Reddy ASN, Golovkin M, editors. Nuclear pre-mRNA Processing in Plants. Berlin, Heidelberg: Springer Berlin Heidelberg; 2008. pp. 83–102. doi:10.1007/978-3-540-76776-3_5

18. Jeong S. SR Proteins: Binders, Regulators, and Connectors of RNA. Mol Cells. 2017/01/26 ed. 2017;40: 1–9. doi:10.14348/molcells.2017.2319

19. Wagner RE, Frye M. Noncanonical functions of the serine-arginine-rich splicing factor (SR) family of proteins in development and disease. BioEssays. 2021;43: 2000242. doi:10.1002/bies.202000242

20. Ji X, Zhou Y, Pandit S, Huang J, Li H, Lin CY, et al. SR Proteins Collaborate with 7SK and Promoter-Associated Nascent RNA to Release Paused Polymerase. Cell. 2013;153: 855–868. doi:10.1016/j.cell.2013.04.028

21. Lin S, Coutinho-Mansfield G, Wang D, Pandit S, Fu X-D. The splicing factor SC35 has an active role in transcriptional elongation. Nat Struct Mol Biol. 2008/07/20 ed. 2008;15: 819–826. doi:10.1038/nsmb.1461

22. Chen T, Cui P, Chen H, Ali S, Zhang S, Xiong L. A KH-Domain RNA-Binding Protein Interacts with FIERY2/CTD Phosphatase-Like 1 and Splicing Factors and Is Important for Pre-mRNA Splicing in Arabidopsis. PLOS Genetics. 2013;9: e1003875. doi:10.1371/journal.pgen.1003875

23. Chong GL, Foo MH, Lin W-D, Wong MM, Verslues PE. Highly ABA-Induced 1 (HAI1)-Interacting protein HIN1 and drought acclimation-enhanced splicing efficiency at intron retention sites. Proc Natl Acad Sci U S A. 2019/10/14 ed. 2019;116: 22376–22385. doi:10.1073/pnas.1906244116

24. Li Y, Guo Q, Liu P, Huang J, Zhang S, Yang G, et al. Dual roles of the serine/arginine-rich splicing factor SR45a in promoting and interacting with nuclear cap-binding complex to modulate the salt-stress response in Arabidopsis. New Phytologist. 2021;n/a. doi:10.1111/nph.17175

25. Albaqami M, Laluk K, Reddy ASN. The Arabidopsis splicing regulator SR45 confers salt tolerance in a splice isoform-dependent manner. Plant Molecular Biology. 2019;100: 379–390. doi:10.1007/s11103-019-00864-4

26. Carvalho RF, Carvalho SD, Duque P. The Plant-Specific SR45 Protein Negatively Regulates Glucose and ABA Signaling during Early Seedling Development in Arabidopsis. Plant Physiol. 2010;154: 772–783. doi:10.1104/pp.110.155523

27. Carvalho RF, Szakonyi D, Simpson CG, Barbosa ICR, Brown JWS, Baena-González E, et al. The Arabidopsis SR45 Splicing Factor, a Negative Regulator of Sugar Signaling, Modulates SNF1-Related Protein Kinase 1 Stability. Plant Cell. 2016;28: 1910–1925. doi:10.1105/tpc.16.00301

28. Xing D, Wang Y, Hamilton M, Ben-Hur A, Reddy ASN. Transcriptome-Wide Identification of RNA Targets of Arabidopsis SERINE/ARGININE-RICH45 Uncovers the Unexpected Roles of This RNA Binding Protein in RNA Processing. Plant Cell. 2015;27: 3294. doi:10.1105/tpc.15.00641

29. Day IS, Golovkin M, Palusa SG, Link A, Ali GS, Thomas J, et al. Interactions of SR45, an SR-like protein, with spliceosomal proteins and an intronic sequence: insights into regulated splicing. The Plant Journal. 2012;71: 936–947. doi:10.1111/j.1365-313X.2012.05042.x

30. Barta A, Kalyna M, Reddy ASN. Implementing a Rational and Consistent Nomenclature for Serine/Arginine-Rich Protein Splicing Factors (SR Proteins) in Plants. Plant Cell. 2010;22: 2926. doi:10.1105/tpc.110.078352

31. Lopato S, Forstner C, Kalyna M, Hilscher J, Langhammer U, Indrapichate K, et al. Network of Interactions of a Novel Plant-specific Arg/Ser-rich Protein, atRSZ33, with atSC35-like Splicing Factors*. Journal of Biological Chemistry. 2002;277: 39989–39998. doi:10.1074/jbc.M206455200

32. Thomas J, Palusa SG, Prasad KVSK, Ali GS, Surabhi G-K, Ben-Hur A, et al. Identification of an intronic splicing regulatory element involved in auto-regulation of alternative splicing of SCL33 pre-mRNA. The Plant journal: for cell and molecular biology. 2012;72: 935–946. doi:10.1111/tpj.12004

33. Yan Q, Xia X, Sun Z, Fang Y. Depletion of Arabidopsis SC35 and SC35-like serine/arginine-rich proteins affects the transcription and splicing of a subset of genes. PLOS Genetics. 2017;13: e1006663. doi:10.1371/journal.pgen.1006663

34. Palusa SG, Ali GS, Reddy ASN. Alternative splicing of pre-mRNAs of Arabidopsis serine/arginine-rich proteins: regulation by hormones and stresses. The Plant Journal. 2007;49: 1091–1107. doi:10.1111/j.1365-313X.2006.03020.x

35. Palusa SG, Reddy ASN. Extensive coupling of alternative splicing of pre-mRNAs of serine/arginine (SR) genes with nonsense-mediated decay. New Phytologist. 2010;185: 83–89. doi:10.1111/j.1469-8137.2009.03065.x

36. Martín G, Márquez Y, Mantica F, Duque P, Irimia M. Alternative splicing landscapes in Arabidopsis thaliana across tissues and stress conditions highlight major functional differences with animals. Genome Biology. 2021;22: 35. doi:10.1186/s13059-020-02258-y

37. Parcy F, Valon C, Raynal M, Gaubier-Comella P, Delseny M, Giraudat J. Regulation of gene expression programs during Arabidopsis seed development: roles of the ABI3 locus and of endogenous abscisic acid. Plant Cell. 1994;6: 1567–1582. doi:10.1105/tpc.6.11.1567

38. Skubacz A, Daszkowska-Golec A, Szarejko I. The Role and Regulation of ABI5 (ABA-Insensitive 5) in Plant Development, Abiotic Stress Responses and Phytohormone Crosstalk. Frontiers in Plant Science. 2016;7: 1884. doi:10.3389/fpls.2016.01884

39. Bensmihen S, Rippa S, Lambert G, Jublot D, Pautot V, Granier F, et al. The homologous ABI5 and EEL transcription factors function antagonistically to fine-tune gene expression during late embryogenesis. Plant Cell. 2002;14: 1391–1403. doi:10.1105/tpc.000869

40. Tian R, Wang F, Zheng Q, Niza VMAGE, Downie AB, Perry SE. Direct and indirect targets of the arabidopsis seed transcription factor ABSCISIC ACID INSENSITIVE3. The Plant Journal. 2020;103: 1679–1694. doi:10.1111/tpj.14854

41. Mao D-D, Tian L-F, Li L-G, Chen J, Deng P-Y, Li D-P, et al. AtMGT7: An Arabidopsis Gene Encoding a Low-Affinity Magnesium Transporter. Journal of Integrative Plant Biology. 2008;50: 1530–1538. doi:10.1111/j.1744-7909.2008.00770.x

42. Surovtseva YV, Shakirov EV, Vespa L, Osbun N, Song X, Shippen DE. Arabidopsis POT1 associates with the telomerase RNP and is required for telomere maintenance. The EMBO Journal. 2007;26: 3653–3661. doi:10.1038/sj.emboj.7601792

43. Tani A, Murata M. Alternative splicing of Pot1 (Protection of telomere)-like genes in Arabidopsis thaliana. Genes & Genetic Systems. 2005;80: 41–48. doi:10.1266/ggs.80.41

44. Kaldis A, Tsementzi D, Tanriverdi O, Vlachonasios KE. Arabidopsis thaliana transcriptional co-activators ADA2b and SGF29a are implicated in salt stress responses. Planta. 2011;233: 749–762. doi:10.1007/s00425-010-1337-0

45. Finkelstein RR, Lynch TJ. The Arabidopsis Abscisic Acid Response Gene <em>ABI5</em> Encodes a Basic Leucine Zipper Transcription Factor. Plant Cell. 2000;12: 599. doi:10.1105/tpc.12.4.599

46. Nishimura N, Yoshida T, Kitahata N, Asami T, Shinozaki K, Hirayama T. ABA-Hypersensitive Germination1 encodes a protein phosphatase 2C, an essential component of abscisic acid signaling in Arabidopsis seed. The Plant Journal. 2007;50: 935–949. doi:10.1111/j.1365-313X.2007.03107.x

47. Nylander M, Svensson J, Palva ET, Welin BV. Stress-induced accumulation and tissue-specific localization of dehydrins in Arabidopsis thaliana. Plant Molecular Biology. 2001;45: 263–279. doi:10.1023/A:1006469128280

48. Yan H, Chaumont N, Gilles JF, Bolte S, Hamant O, Bailly C. Microtubule self-organisation during seed germination in Arabidopsis. BMC Biol. 2020;18: 44–44. doi:10.1186/s12915-020-00774-8

49. Costa MCD, Righetti K, Nijveen H, Yazdanpanah F, Ligterink W, Buitink J, et al. A gene co-expression network predicts functional genes controlling the re-establishment of desiccation tolerance in germinated Arabidopsis thaliana seeds. Planta. 2015/03/26 ed. 2015;242: 435–449. doi:10.1007/s00425-015-2283-7

50. Moore R, Smith JD. Growth, graviresponsiveness and abscisic-acid content of Zea mays seedlings treated with Fluridone. Planta. 1984;162: 342–344. doi:10.1007/BF00396746

51. Ullah H, Chen J-G, Wang S, Jones AM. Role of a Heterotrimeric G Protein in Regulation of Arabidopsis Seed Germination. Plant Physiology. 2002;129: 897–907. doi:10.1104/pp.005017

52. Lin P-C, Hwang S-G, Endo A, Okamoto M, Koshiba T, Cheng W-H. Ectopic Expression of ABSCISIC ACID 2/GLUCOSE INSENSITIVE 1 in Arabidopsis Promotes Seed Dormancy and Stress Tolerance. Plant Physiology. 2007;143: 745–758. doi:10.1104/pp.106.084103

53. Schwartz SH, Leon-Kloosterziel KM, Koornneef M, Zeevaart JAD. Biochemical Characterization of the aba2 and aba3 Mutants in Arabidopsis thaliana. Plant Physiology. 1997;114: 161–166. doi:10.1104/pp.114.1.161

54. Finkelstein RR, Li Wang M, Lynch TJ, Rao S, Goodman HM. The Arabidopsis Abscisic Acid Response Locus <em>ABI4</em> Encodes an APETALA2 Domain Protein. Plant Cell. 1998;10: 1043. doi:10.1105/tpc.10.6.1043

55. Söderman EM, Brocard IM, Lynch TJ, Finkelstein RR. Regulation and Function of the Arabidopsis ABA-insensitive4 Gene in Seed and Abscisic Acid Response Signaling Networks1. Plant Physiology. 2000;124: 1752–1765. doi:10.1104/pp.124.4.1752

56. Laby RJ, Kincaid MS, Kim D, Gibson SI. The Arabidopsis sugar-insensitive mutants sis4 and sis5 are defective in abscisic acid synthesis and response. The Plant Journal. 2000;23: 587–596. doi:10.1046/j.1365-313x.2000.00833.x

57. Finkelstein RR. Mutations at two new Arabidopsis ABA response loci are similar to the abi3 mutations. The Plant Journal. 1994;5: 765–771. doi:10.1046/j.1365-313X.1994.5060765.x

58. Li Y, Zheng L, Corke F, Smith C, Bevan MW. Control of final seed and organ size by the DA1 gene family in Arabidopsis thaliana. Genes & Development. 2008;22: 1331–1336. doi:10.1101/gad.463608

59. Lopez-Molina L, Mongrand S, McLachlin DT, Chait BT, Chua N-H. ABI5 acts downstream of ABI3 to execute an ABA-dependent growth arrest during germination. The Plant Journal. 2002;32: 317–328. doi:10.1046/j.1365-313X.2002.01430.x

60. Lopez-Molina L, Mongrand S, Chua NH. A postgermination developmental arrest checkpoint is mediated by abscisic acid and requires the ABI5 transcription factor in Arabidopsis. Proc Natl Acad Sci U S A. 2001/04/03 ed. 2001;98: 4782–4787. doi:10.1073/pnas.081594298

61. Ali GS, Golovkin M, Reddy ASN. Nuclear localization and in vivo dynamics of a plant-specific serine/arginine-rich protein. The Plant Journal. 2003;36: 883–893. doi:10.1046/j.1365-313X.2003.01932.x

62. Tillemans V, Leponce I, Rausin G, Dispa L, Motte P. Insights into Nuclear Organization in Plants as Revealed by the Dynamic Distribution of <em>Arabidopsis</em> SR Splicing Factors. Plant Cell. 2006;18: 3218. doi:10.1105/tpc.106.044529

63. Rausin G, Tillemans V, Stankovic N, Hanikenne M, Motte P. Dynamic Nucleocytoplasmic Shuttling of an Arabidopsis SR Splicing Factor: Role of the RNA-Binding Domains. Plant Physiology. 2010;153: 273–284. doi:10.1104/pp.110.154740

64. Wang P, Xue L, Batelli G, Lee S, Hou Y-J, Van Oosten MJ, et al. Quantitative phosphoproteomics identifies SnRK2 protein kinase substrates and reveals the effectors of abscisic acid action. Proc Natl Acad Sci USA. 2013;110: 11205–11210. doi:10.1073/pnas.1308974110

65. Umezawa T, Sugiyama N, Takahashi F, Anderson JC, Ishihama Y, Peck SC, et al. Genetics and Phosphoproteomics Reveal a Protein Phosphorylation Network in the Abscisic Acid Signaling Pathway in Arabidopsis thaliana. Sci Signal. 2013;6: rs8. doi:10.1126/scisignal.2003509

66. de la Fuente van Bentem S, Anrather D, Dohnal I, Roitinger E, Csaszar E, Joore J, et al. Site-Specific Phosphorylation Profiling of Arabidopsis Proteins by Mass Spectrometry and Peptide Chip Analysis. J Proteome Res. 2008;7: 2458–2470. doi:10.1021/pr8000173

67. Howard JM, Sanford JR. The RNAissance family: SR proteins as multifaceted regulators of gene expression. Wiley Interdiscip Rev RNA. 2014/08/22 ed. 2015;6: 93–110. doi:10.1002/wrna.1260

68. Clough SJ, Bent AF. Floral dip: a simplified method for Agrobacterium -mediated transformation of Arabidopsis thaliana. The Plant Journal. 1998;16: 735–743. doi:10.1046/j.1365-313x.1998.00343.x

69. Karimi M, Inzé D, Depicker A. GATEWAY^TM^ vectors for Agrobacterium-mediated plant transformation. Trends in Plant Science. 2002;7: 193–195. doi:10.1016/S1360-1385(02)02251-3

70. Meyer K, Stecca KL, Ewell-Hicks K, Allen SM, Everard JD. Oil and protein accumulation in developing seeds is influenced by the expression of a cytosolic pyrophosphatase in Arabidopsis. Plant Physiol. 2012/05/07 ed. 2012;159: 1221–1234. doi:10.1104/pp.112.198309

71. Sundaresan V, Springer P, Volpe T, Haward S, Jones JD, Dean C, et al. Patterns of gene action in plant development revealed by enhancer trap and gene trap transposable elements. Genes & Development. 1995;9: 1797–1810. doi:10.1101/gad.9.14.1797

72. Carvalho SD, Saraiva R, Maia TM, Abreu IA, Duque P. XBAT35, a Novel Arabidopsis RING E3 Ligase Exhibiting Dual Targeting of Its Splice Isoforms, Is Involved in Ethylene-Mediated Regulation of Apical Hook Curvature. Molecular Plant. 2012;5: 1295–1309. doi:10.1093/mp/sss048

73. Tapial J, Ha KCH, Sterne-Weiler T, Gohr A, Braunschweig U, Hermoso-Pulido A, et al. An atlas of alternative splicing profiles and functional associations reveals new regulatory programs and genes that simultaneously express multiple major isoforms. Genome Res. 2017/08/30 ed. 2017;27: 1759–1768. doi:10.1101/gr.220962.117

74. Braunschweig U, Barbosa-Morais NL, Pan Q, Nachman EN, Alipanahi B, Gonatopoulos-Pournatzis T, et al. Widespread intron retention in mammals functionally tunes transcriptomes. Genome Res. 2014/09/25 ed. 2014;24: 1774–1786. doi:10.1101/gr.177790.114

75. Huang DW, Sherman BT, Lempicki RA. Systematic and integrative analysis of large gene lists using DAVID bioinformatics resources. Nature Protocols. 2009;4: 44–57. doi:10.1038/nprot.2008.211

